# Vitamin and amino acid auxotrophy in anaerobic consortia operating under methanogenic condition

**DOI:** 10.1101/128660

**Authors:** Valerie Hubalek, Moritz Buck, BoonFei Tan, Julia Foght, Annelie Wendeberg, David Berry, Stefan Bertilsson, Alexander Eiler

**Affiliations:** Department of Ecology and Genetics, Limnology and Science for Life Laboratory, Uppsala University, Sweden; NBIS, National Bioinformatics Infrastructure Sweden, Uppsala, Sweden; Department of Biological Sciences, University of Alberta, Edmonton, Alberta, Canada; Center for Environmental Sensing and Modeling, Singapore-MIT Alliance for Research and Technology; Centre for Environmental Research, Environmental Microbiology, Leipzig, Germany; Department of Microbiology and Ecosystem Science, Division of Microbial Ecology, University of Vienna, Vienna, Austria; eDNA solutions AB, Mölndal, Sweden

## Abstract

Syntrophy among Archaea and Bacteria facilitates the anaerobic degradation of organic compounds to CH_4_ and CO_2_. Particularly during aliphatic and aromatic hydrocarbon mineralization, as in crude oil reservoirs and petroleum-contaminated sediments, metabolic interactions between obligate mutualistic microbial partners are of central importance^1^. Using micro-manipulation combined with shotgun metagenomic approaches, we disentangled the genomes of complex consortia inside a short chain alkane-degrading cultures operating under methanogenic conditions. Metabolic reconstruction revealed that only a small fraction of genes in the metagenome-assembled genomes of this study, encode the capacity for fermentation of alkanes facilitated by energy conservation linked to H_2_ metabolism. Instead, inferred lifestyles based on scavenging anabolic products and intermediate fermentation products derived from detrital biomass was a common feature in the consortia. Additionally, inferred auxotrophy for vitamins and amino acids suggests that the hydrocarbon-degrading microbial assemblages are structured and maintained by multiple interactions beyond the canonical H_2_-producing and syntrophic alkane degrader–methanogen partnership^2^. Our study uncovers the complexity of ‘interactomes’ within microbial consortia mediating hydrocarbon transformation under anaerobic conditions.

---

Microbial consortia that degrade hydrocarbons under anaerobic conditions hold the potential to influence hydrocarbon profiles with either adverse or beneficial effects on the recovery and refining of oil^3–5^. In environmental restoration, microbial degradation is seen as an effective and environmental-friendly treatment of oil-contaminated sites. Still, in anaerobic systems, bioremediation efforts are challenged by low efficiency and poor success. Additionally there is a severe problem with unwanted contamination of adjacent systems with pollutants such as toxic metals, naphthenic acids and polycyclic aromatic compounds^6–8^ as well as accelerated release of the potent greenhouse gas methane from the biodegradation process^9^.

Analyses of chemical profiles and metagenomic data from hydrocarbon-degrading consortia maintained in laboratory-scale cultures^10–15^ have identified some of the organisms and the metabolic pathways involved in the anaerobic degradation of various hydrocarbons, such as alkanes, benzenes and naphthalenes^14,19-21,16,17^. The anaerobic degradation of hydrocarbons is usually a two-to three-step process involving fermentation either to simple substrates for methanogenesis (hydrogen, formate, acetate, CO_2_) or to other metabolites (e.g. lactate, ethanol, propionate, butyrate, fatty acids) that can undergo secondary fermentation yielding aforementioned simple methanogenic substrates^18–20^.

Still, we are far from having a detailed understanding of the molecular mechanisms that establish and maintain such syntrophic methanogenic consortia where methanogens deplete the partial pressure of hydrogen facilitating the fermentation process by making it thermodynamically favourable. Additional members of the consortium that scavenge metabolic byproducts and thereby participate in the biodegradation process as secondary consumers, have thus far been overlooked^21^ and their additional functions in the consortia remain hidden. One possible role of such scavengers could be as providers of essential nutrients to salvage other consortia members from auxotrophies while they also alter the thermodynamic equilibrium. It previously has been suggested that inability of individual community members to synthesize the full range of organic nutrients required for growth may be a favourable trait for communities and consortia operating under energy-limited conditions^1^. Reconstructing metabolic pathways and assigning functions among community members using genomics is a first step towards a more holistic understanding of microbial consortia mediating anaerobic degradation of hydrocarbons^2^.

For this study, we extracted high-quality genomes of syntrophic bacteria from public databases. To identify taxa that have the potential to grow syntrophically, we used gene annotations predicting enzymes involved in reverse and direct electron transfer. Many syntrophic bacteria are known to rely on the capacity to perform reverse electron transport-driven energy-conserving H_2_ production through electron-confurcating hydrogenase (H_2_ase) in combination with reduced ferredoxin^21^. Our analysis confirmed a wide distribution of membrane-bound, ion-translocating ferredoxin:NAD+ oxidoreductase and confurcating hydrogenase that could directly couple the oxidation of NADH and reduced ferredoxin to produce hydrogen (Suppl. Tab. 1). Additionally, most of the putatively syntrophic bacterial isolates lacked important genes involved in biosynthesis of one or more amino acids and vitamins, indicating that they rely on precursors (or the co-factors) provided by other bacteria to produce fully functional enzymes with required cofactors and prosthetic groups (Suppl. Fig. 1).

Next, we attempted to resolve these interactions beyond the available genomes in public databases by isolating natural methanogenic consortia. Short-chain alkane degrading enrichment cultures (SCADCs) from mature fine tailings of the Mildred Lake Settling Basin^13^ were used to obtain 13 shotgun metagenomes from various subcultures of the original enrichment culture (for details on the metagenomes see suppl. Tab. 2). The cultures have previously been described in several metagenomic studies inferring the broader metabolic roles and identity of the community members at the phylum level^13^ (for taxonomic composition see Suppl. Fig. 2). These included eight samples where each individual sample contained approximately ten individually picked filaments, pooled based on the presence or absence of visible epibionts. The physical manipulation for sequencing was an attempt to isolate members of the consortia that exchange co-factors or their precursors in close physical contact such as syntrophs, versus free-living community members with peripheral roles in alkane degradation.

From the pool of ∼80 selected filaments, we obtained a metagenome co-assembly of 805606 scaffolds totaling ∼1 Gbp with an N50-length of 4902 bp using MEGAHIT. Downstream coverage and nucleotide composition-based binning of the 150698 scaffolds of at least 1000 bp length, resulted in 307 bins with 45 matching our criteria for being defined as high-quality metagenome assembled genomes (MAGs). As indicated by detailed annotations of these MAGs using the ‘ MicroScope’ pipeline^22^, the most likely hydrocarbon degraders in the culture are putative syntrophic bacteria related to *Syntrophus*, *Pelotomaculum*, *Desulfobacterium* and *Clostridium*, all having the potential for n-alkane activation, betaoxidation and fumarate regeneration. The genes indicative for these pathways were alkylsuccinate synthase and benzylsuccinate synthase (C-H activation), methylmalonyl CoA mutase (carbon skeleton rearrangement) and methylmalonyl-CoA decarboxylase (decarboxylation) for n-alkane activation; enoyl-CoA hydratase, β-hydroxyacyl-CoA dehydrogenase and acyl-CoA acetyl-transferase for beta-oxidation; propionyl-CoA carboxylase, methylmalonyl-CoA mutase, succinyl-CoA synthetase and succinate dehydrogenase for fumarate regeneration^14^. Most genomes encoded enzymes associated with fermentation and coupled pathways for acetogenesis (i.e. reductive acetyl-CoA or Wood-Ljungdahl pathway), a process by which acetate is produced from CO_2_ and an electron donor (Figure 1). The five methanogen genomes reconstructed from the contig bins belonged to the genera *Methanoculleus* and *Methanosaeta* in the class *Methanomicrobia* and to the class *Thermoplasmata* As previously described, *Methanosaeta* represent the acetoclastic methanogens while *Methanoculleus* represent hydrogenotrophic methanogens, and *Thermoplasmata* represent methylotrophic methanogens^23^. Besides these differences in substrates used for methane production, they all encode complete sets of nitrogen fixation genes. For detailed exploration of the genomes we direct the reader to the online sources of the ‘ MicroScope’ pipeline^22^ where each genome’s individual KEGG and biocyc maps can be inspected.

**Figure 1.**
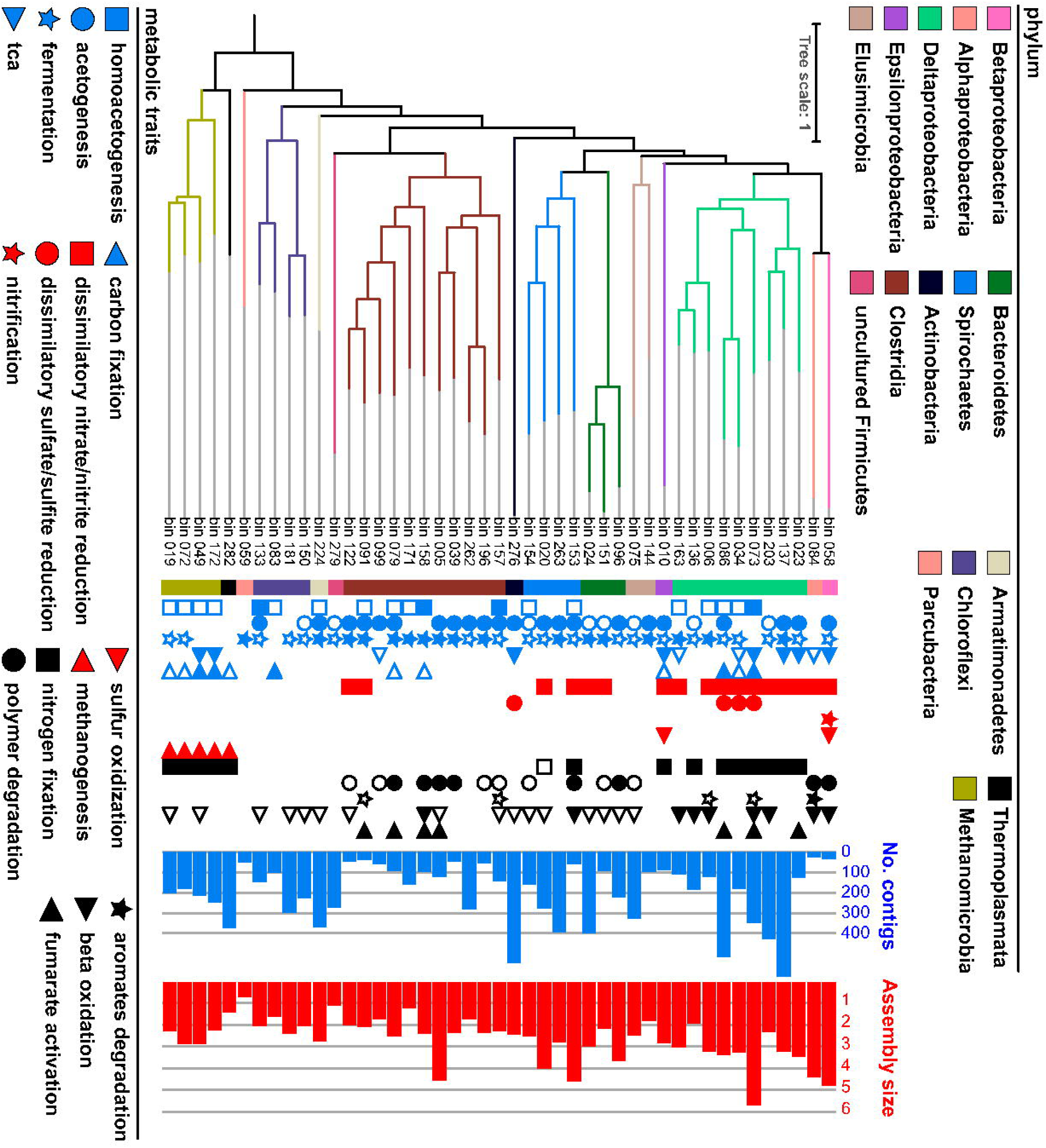
A phylogenomic tree of recovered genomes as computed by PhyloPhlAn^41^. The colorstrip delineates taxonomic affiliation. The symbols show the metabolic traits, based on genome annotations. Traits with high support are indicated by closed symbols, while traits with low support have open symbols, and categories with missing symbols indicate that no indications of the trait were found. The two barcharts on the far right show the number of contigs assigned to each bin and the assembly size (in Mbp) contained in the recovered genome bins.

In addition to classical syntrophic interactions described in hydrocarbon-degrading communities, i.e. where one partner produces metabolites such as H_2_, acetate, ethanol and/or small organic intermediates, which are then used by methanogens and secondary degraders, our genomic reconstructions point to additional metabolic dependencies. Firstly, the scavenging of metabolic products derived from detrital biomass is indicated by the presence of genes encoding transporters for a wide diversity of proteins and fatty acids, as well as genes involved in the degradation of intermediate fermentation products (i.e. C1 compounds and alcohols) (Figure 2). Genomes of these secondary degraders belonged to the deltaproteobacterial genera *Desulfovibrio*, *Desulfomicrobium* and *Desulfobulbus* previously reported to use H_2_, organic acids, formate, or short-chain alcohols as electron donors for sulfate reduction^24–26^. In addition to their potential to grow by using anaerobic respiration, these genera possessed genes associated with fermentation of a wide range of alcohols and organic acids such as lactate, amino and carboxylic acids.

**Figure 2:**
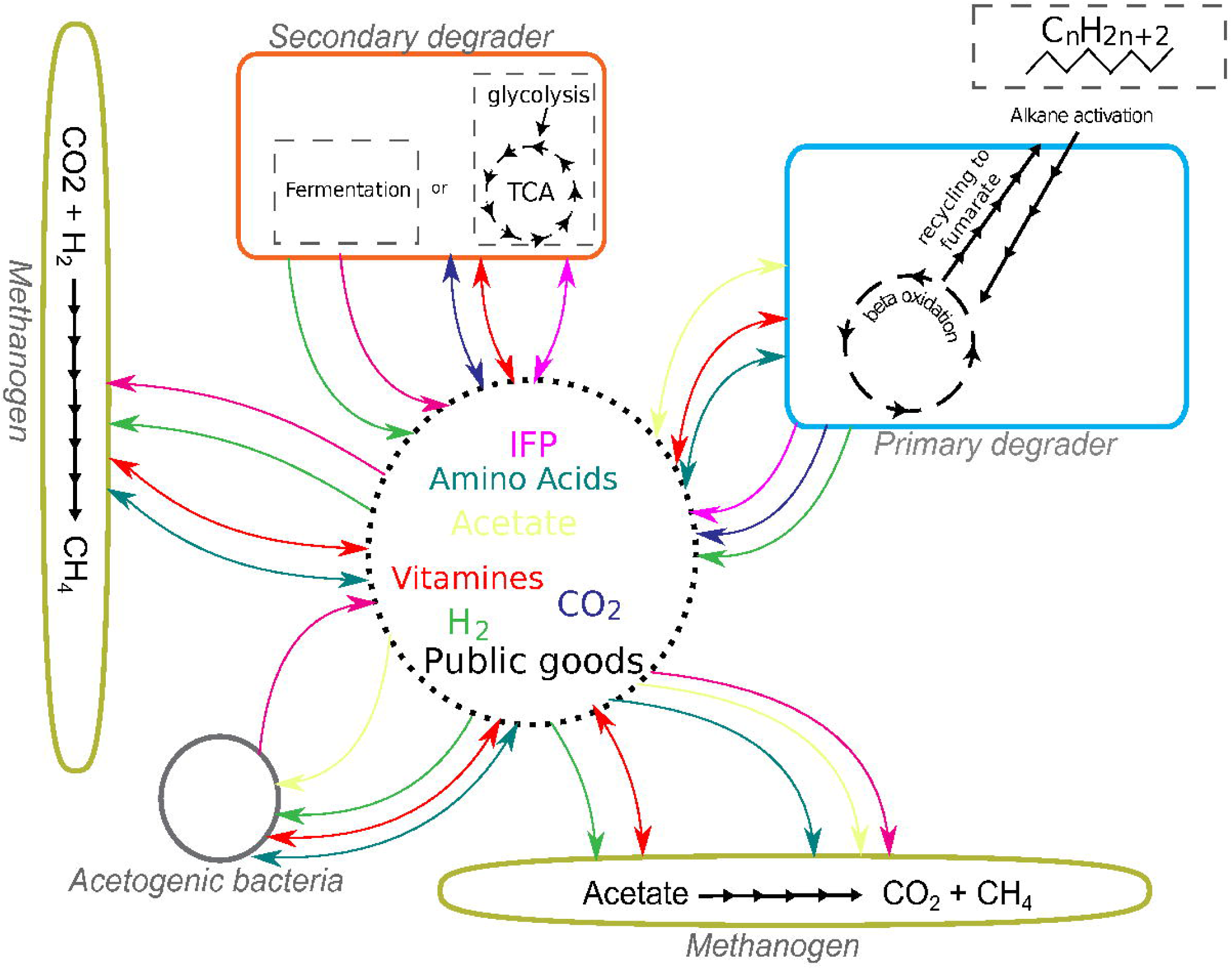
Model of potential metabolic pathways and metabolic cross-feeding in the short-chain alkane degrading culture (SCADC) representing (green) archaeal filaments, (blue) alkane degraders, (red) intermediate metabolite scavengers and secondary degraders, (grey) acetogenic bacteria. In addition, the figure shows the inferred syntrophic interactions from hydrogen (green), carbon dioxide (dark blue) and acetate (green) to intermediate fermentation metabolites (IFMs; violet) transfer as well as other forms of metabolic interdependencies such as cross-feeding of costly metabolites such as vitamins (red) and amino acids (black).

Like many of their cultivated and genome sequenced relatives, the MAGs from the putative sulfate reducers in these consortia revealed auxotrophies for vitamins and amino acids (Figure 3). Even though the absence of pathways, defined by less than half of the genes present in the respective pathways, need to be assessed with caution because of the partial nature of the individual MAGs, analogous auxotrophies in closely related MAGs and closed genomes from public databases provide strong support for widespread auxotrophy. Examples of essential metabolite pathways whose biosynthetic gene distribution is patchy in the SCADC MAGs are those for biosynthesis of vitamins B6 (pyridoxal), B12 (adenosylcobalamin), and H (biotin). Earlier large-scale genome sequence analyses have previously suggested that corrinoids, essential cofactors for vitamins^27^, are widely shared by over 70% of the sequenced bacterial species, while less than 25% of species synthesize corrinoids *de novo*^28,29^. The existence of over a dozen structurally distinct corrinoids produced and used by various Bacteria and Archaea^30^ can thus hold a central role in dictating microbial community assembly and functioning, as it is clear that the structure of the precursor, which can be a benzimidazole, purine or a phenolic compound^30^, influences the degree to which a particular corrinoid can be used by a given organism^31,32^.

**Figure 3:**
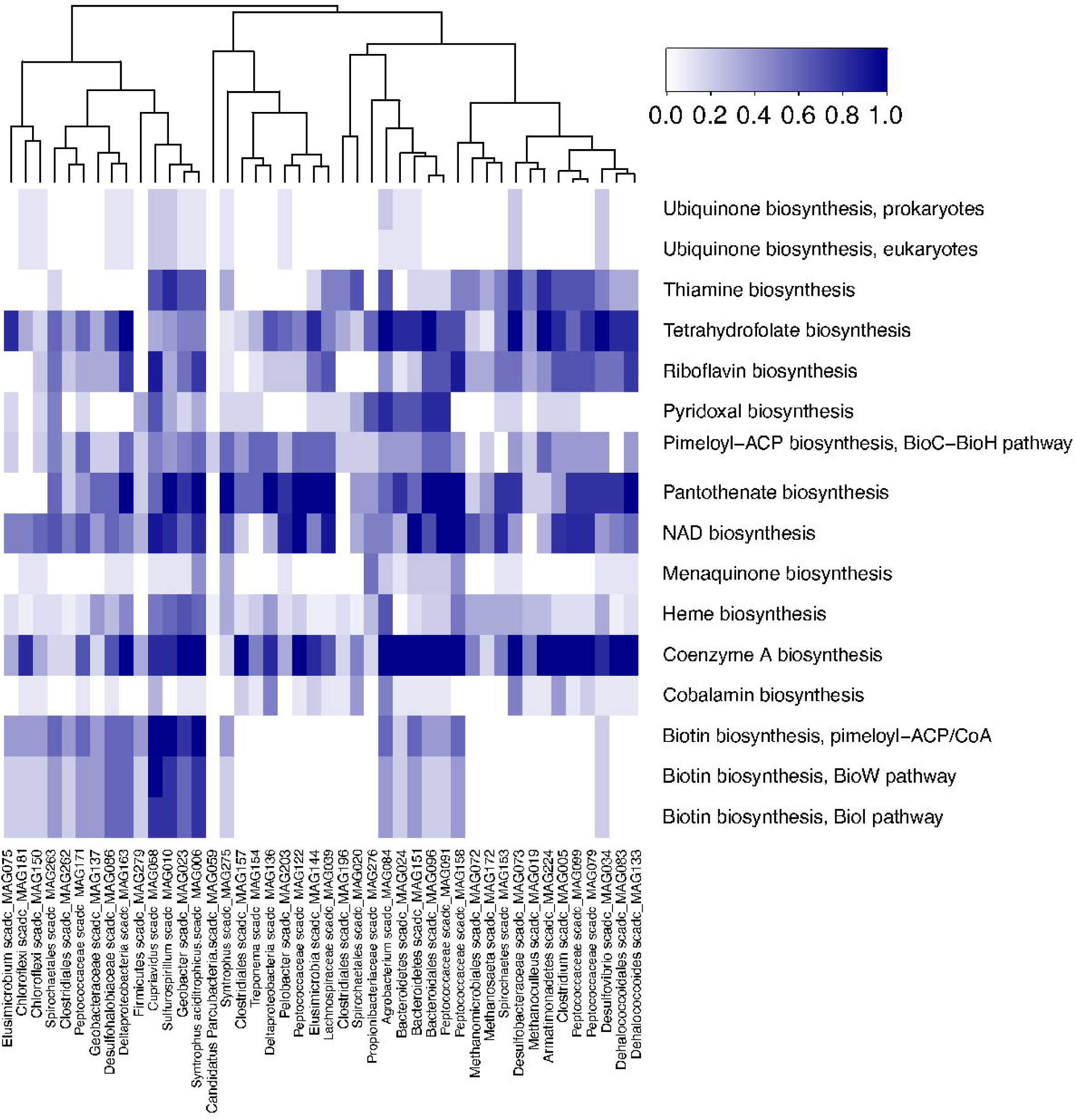
Heat map of amino acid (A) and vitamin (B) biosynthetic capabilities of taxonomic bins obtained from the short-chain alkane degrading culture (SCADC). Prototrophy predictions for each amino acid and vitamin are based on ‘pathway completion’ value i.e, the number of reactions for pathway x in a given organism/total number of reactions in the same pathway x defined in the MetaCyc or KEGG databases. Value of 1/0 indicates all/none of the key enzymes involved in the biosynthesis. Dendograms represent clustering based on potential biosynthesis profiles. To verify widespread auxotrophy for vitamins and amino acids in syntrophic prokaryotes, a similar analysis was performed using genomes available from genera representing our taxonomic bins (for details see Supplementary Figure 1). As a definition of absence we used a threshold of less than 0.5 pathway completeness.

Likewise, the absence of genes underlying biosynthesis of essential amino acids is apparent in particular taxonomic groups (Figure 3B). The loss of essential biosynthesis function in various taxonomic groups has previously been associated with energy- and nutrient-limitation selecting for small cells with reduced genomes such as found in the open ocean and deep biosphere^33,34^. Particularly costly functions are absent in most members of the SCADC community, resulting in a community critically depending on essential metabolite providers. Biosynthesis of certain vitamins such as adenosylcobalamin and specific amino acids such as phenylalanine, tyrosine and tryptophan, is energetically costly and require a large set of genes. This loss of function and subsequent adaptations to a cooperative lifestyle under nutrient- and energy-limitation forms the foundation for the Black Queen hypothesis^35^.

In the context of the SCADC, the Black Queen hypothesis implies that, in addition to fermentation products disposal by methanogens, other types of “public goods” can be exchanged. Such "public goods" are provided by a few taxa hosting the enzymes to perform this energetically costly biosynthesis while benefitting the other organisms in their vicinity. This enhanced cross feeding is logical if one considers the need for metabolic processes to be optimized for reduced metabolic burden at the level of an individual cell living at the thermodynamic limit.

To conclude, we show that the interactome of syntrophic alkane degrader–methanogen partnerships surpass the standard electron- and simple carbon compound-transfer. Biosynthesis of vitamins and other energy-expensive metabolites seem to be unequally distributed among community members and we propose that this facilitate energy conservation at the thermodynamic limit. It can be further speculated that vitamin and amino acid augmentations could possibly enhance petroleum bioaugmentation processes in remediation of hydrocarbon-contaminated sites.

## Methods

### Culturing

An enriched short-chain alkane-degrading culture (SCADC) was enriched from mature fine tailings of the Mildred Lake Settlings Basin as detailed by Tan *et al.*^13^. In short, the primary enrichment cultures were established in multiple replicates by adding 25 ml oil sand tailings to 50 ml of mineral methanogenic medium^36^, under a headspace of O_2_-free 30% CO_2_–N_2_ balance gas. After an initial 1-year incubation in the dark at room temperature, the primary cultures were pooled and used to inoculate multiple replicates of first transfer cultures into fresh methanogenic medium. After a second 6-month incubation, these cultures were pooled as an inoculum for the experimental SCADC enrichment cultures, where 0.1% (v/v) of an alkane mixture comprising equal volumes of C_6_-C_9_ compounds was added to the community and incubated at least 4 months at 28 °C in the dark, prior to further subculturing^13^. The samples analyzed in the present study were from several transfers of the parent SCADC culture. Three bottles containing 75-ml cultures were each sampled three times using micromanipulation, originally to capture filamentous bacteria seen to carry attached smaller cocci (Fig. 1). Fluorescence in situ hybridization (FISH) using probes MX825b and MX825c^37^ revealed that most filaments represent *Methanosaeta* with associated bacterial cells.

### Filament sampling

Micromanipulation was performed using an Eppendorf TransferMan® NK 2 at the Department of Microbial Ecology, University of Vienna. Each slide used for micromanipulation was prepared by cleaning with “DNA AWAY” surface decontaminant (ThermoFisher Scientific) and rinsed in sterile phosphate-buffered saline. Filaments were picked from a 12 μl culture subsample by using microscopy (Axio Observer.D1, Zeiss) at 32x magnification, captured in a 4 μm diameter capillary (TransferTip®, Eppendorf) and subsequently transferred into a sterile tube. Approximately 10 filaments were collected from each of eight cultures and pooled according to coherent morphology.

Samples with pooled filaments (n = 8) were then lysed by subjecting them to four freeze-thaw cycles, freezing at -80°C followed by heat treatment at 70°C for 5 minutes. These samples were subjected to whole genome amplification with multiple displacement amplification (MDA) using the REPLI-g® Mini kit according to the manufacturer’s instructions (Qiagen).

### Whole culture DNA isolation

From the SCADC culture, 100 ml were filtered through 0.2 μm Supor® Membrane Disc Filters (Pall). Prior to bead beating, the above mentioned freeze-thaw cell lysis was conducted, as preceding tests showed that the filaments needed a more vigorous treatment. The DNA was then isolated from the filters by bead-beating using the PowerSoil® DNA Isolation Kit (MoBio) according to the manufacturer’s instructions, and sequenced as described below. No whole genome amplification with multiple displacement amplification (MDA) was performed on these samples.

### Library preparation and sequencing

The SciLifeLab SNP/SEQ sequencing facility at Uppsala University performed the library preparation and the sequencing. First, 10 ng of genomic DNA was sheared using a focused ultrasonicator (Covaris E220). The sequencing libraries were prepared with the Thruplex FD Prep kit (Rubicon Genomics) according to manufacturer’s protocols (R40048-08, QAM-094-002). Library size selection was made using AMPpure XP beads (Beckman Coulter) in 1:1 ratio with the DNA sample volume. The prepared sample libraries were quantified using KAPA Biosystems next-generation sequencing library qPCR kit and analysing using a StepOnePlus (Life Technologies) real-time PCR instrument. The quantified libraries were then prepared for sequencing on the Illumina HiSeq sequencing platform with a TruSeq paired-end cluster kit, v3, and Illumina’s cBot instrument to generate a clustered flowcell for sequencing. Sequencing of the flowcell contents was performed using an Illumina HiSeq2500 sequencer with Illumina TruSeq SBS sequencing kit v3, following a 2x100 indexed high-output run recipe.

### Assembly and binning of metagenome reads

Reads were filtered based on their quality scores using Sickle (version 1.210)^38^. The read-set was digitally normalized using khmer (arXiv:1203.4802) and subsequently assembled with MEGAHIT^39^. Contigs were scaffolded with the BESST software^40^. Coverage was computed by mapping back the reads to the scaffolds using bbmap (BBMap - Bushnell B. - sourceforge.net/projects/bbmap/) and for computing coverage, bedtools (version 2.18.2)^41^ was used. Scaffolds were binned using MetaBAT (doi:10.7717/peerj.1165) The criteria for high quality bins, so-called MAGs, are based on CheckM(doi:10.1101/gr.186072.114) estimates of completeness (>70%) and contamination (<5%).

### Processing of mined data

In addition to data acquired from the newly sequenced samples, we also accessed three datasets from the same original SCADC, downloaded from the NCBI Short Read Archive (SRA): SRR636559 (a high depth HiSEQ dataset), and SRR634694 and SRR634695 (two Roche 454 datasets). These archival sequences were treated bioinformatically in the same way as the new sequences and co-assembled with them.

### Annotation

Initial phylogenetic attribution of the MAGs was performed using PhyloPhlAn^42^ on proteins predicted with Prokka (version 1.9)^43^, supplemented with a custom set of MAGs, single amplified genomes (SAGs) and whole genome sequences accessed from JGI IMG/M. PFAM annotations were performed using the Pfam database 27.0^44^ and HMMER^45^ (version 3.1b1) with default parameters. Detailed genome annotation analyses were performed using the ‘ MicroScope’ pipeline^22^ with automatic annotations assisted by manual curation as described in the integrated bioinformatics tools and the proposed annotation rules. Reconstructed MAGs are deposited on MicroScope platform^22^ with identifier ‘ scadc_MAG’.

## Acknowledgements

We acknowledge the Uppsala Multidisciplinary Center for Advanced Computational Science (UPPMAX) for access to data storage and computing resources under project b2013274. This work was supported by the Swedish Research Council VR (Grant 2012-4592 to AE and 2012-3892 to SB), the Swedish Foundation for Strategic Research (Grant ICA010-015 to AE), Swedish Research Council Formas (project 2012-986 to SB) and the Helmholtz Initiative and Networking Fund.

## Supplementary Material

**Supplementary Figure 1.**
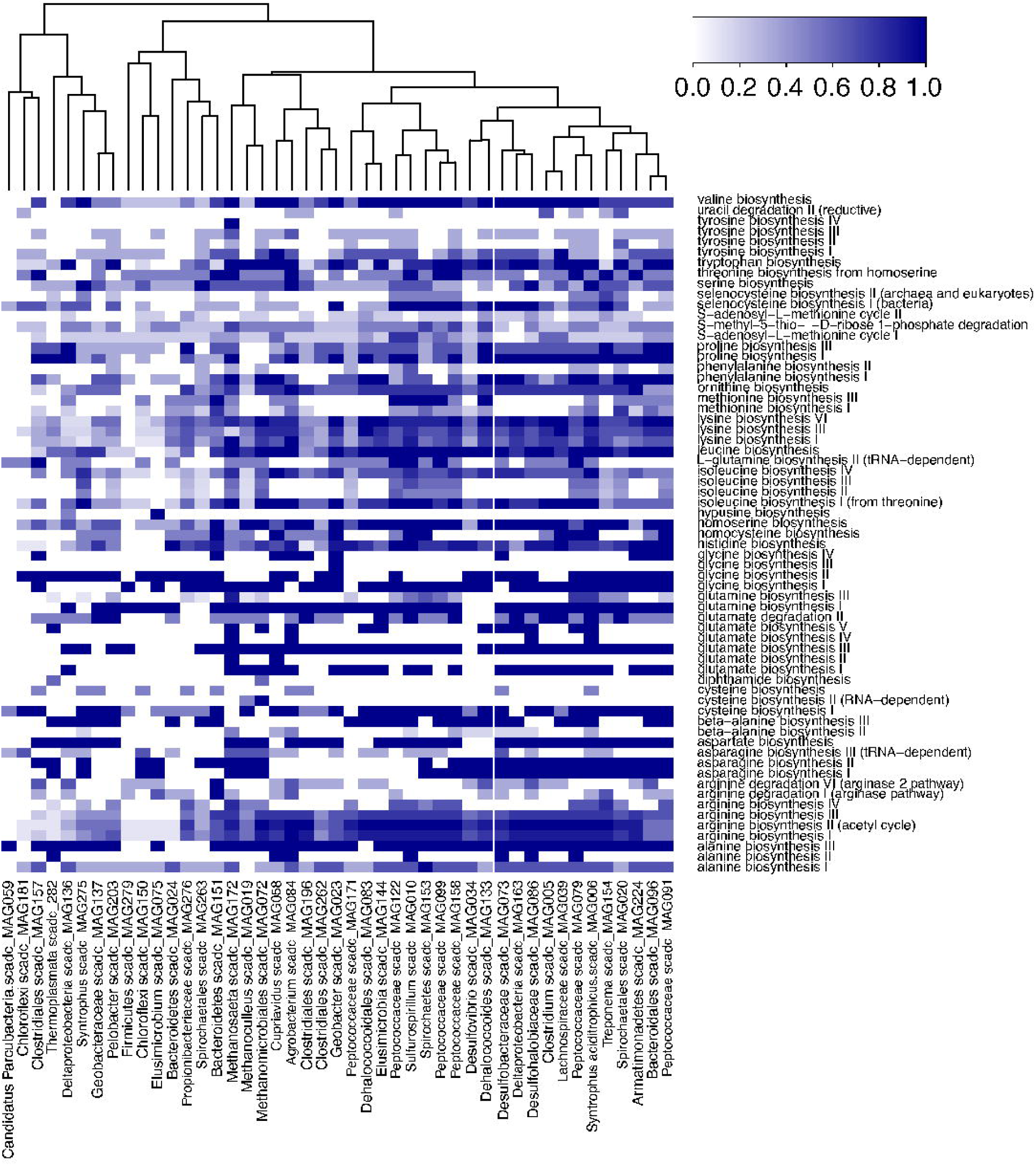
Heat map of amino acid (A) and vitamin (B) biosynthetic capabilities of selected genomes obtained from MaGe. Prototrophy predictions for each amino acid and vitamin are based on ‘pathway completion’ value i.e, the number of reactions for pathway x in a given organism/total number of reactions in the same pathway x defined in the MetaCyc or KEGG databases. Value of 1/0 indicates all/none of the key enzymes involved in the biosynthesis. Dendograms represent clustering based on potential biosynthesis profiles. This resembles wide spread auxotrophy for vitamins and amino acids as inferred for the short-chain alkene degrading culture (SCADC).

Supplementary Figure 2. Taxonomic composition of the SCADC in the different samples used for shotgun metagenomic sequencing. The 16S rRNA gene Data was produced following the procedure as outlined in Sinclair *et al*. (2015).

Supplementary Table 1. Distribution of membrane-bound, ion-translocating ferredoxin:NAD+ oxidoreductase and confurcating hydrogenase that could directly couple the oxidation of NADH and reduced ferredoxin to produce hydrogen.

Supplementary Table 2. Source of metagenome and summary of sequencing data. “+Epibiont” indicates samples including filaments with epibioints whereas “-Epibioints” without epibionts.

